# Comparative transcriptomics unveil distinctive metabolic pathway of phosphonate utilization by diatom *Phaeodactylum tricornutum*

**DOI:** 10.1101/2022.02.09.479573

**Authors:** Huilin Shu, Yanchun You, Hongwei Wang, Jingtian Wang, Ling Li, Xin Lin, Jian Ma

**Author notes:** Xin Lin; 05922186779. Jian Ma; 05922186916.

## Abstract

Phosphonates are important constituents of marine organic phosphorus, however, the bioavailability and catabolism of phosphonates by eukaryotic phytoplankton remain enigmatic. Here, we use diatom *Phaeodactylum tricornutum* to investigate the bioavailability of phosphonates and elaborate the underlying molecular mechanism. Our results showed that 2-aminoethylphosphonic (2-AEP) can be utilized as alternative phosphorus source. Comparative transcriptomics unveil the 2-AEP utilization comprising two steps, molecular uptake through clathrin-mediated endocytosis and incorporation into the membrane phospholipids in the form of diacylglyceryl-2-AEP (DAG-2-AEP). In the global ocean, we found the prevalence of key genes responsible for vesicle formation (*CLTC, AP-2*) and DAG-AEP synthesis (*PCYT2* and *EPT1*) in diatom assemblage. In accordance with the observation of elevated transcript abundance in cold waters, our culture experiments showed that cells grown in 2-AEP are more active at lower temperature. This study elucidated a distinctive mechanism of phosphonate utilization by diatom and inspected the ecological implications in adaptive mechanism.

## Introduction

Phosphorus (P) is an essential element for living organisms. Dissolved inorganic phosphate (DIP) is the preferable form to microorganisms, but is scarce and often limiting in the surface ocean [1]. The dissolved organic phosphorus (DOP) has been recognized as a vital alternative P source and the bioavailability of different forms of DOPs have arisen many discussions [2, 3]. Phosphonates are a broad family of organic phosphorus that contains C-P bond, estimated to contribute 25% of the total DOP [4] and play an important role in the P redox cycle [5]. C-P bond is much more stable in comparison with C-O-P bond in phosphate esters, because it is resistant to chemical hydrolysis, thermal decomposition and photolytic degradation [6].

Phosphonates are largely found as biogenic compounds synthesized by a wide range of organisms either in free state or combined with proteins [7], lipids [8] and glycans [9] in lower organisms. 2-aminoethylphosphonic (2-AEP) is the first identified natural phosphonate and one of the most abundant and ubiquitous phosphonates in the natural environment [6, 10], suggested protecting cells against predators while incorporated into membrane phospholipids [11]. For example, glycerophospholipid DAG-2-AEP [6], being a constituent in membrane phospholipids, is considered to protect the cells from enzymatic degradation or increase the structural rigidity attributed to the stability of C-P bond [12]. Besides that, current evidence indicates that biogenic phosphonates include 2-AEP have a significant and previously underestimated role in marine P cycle [13].

Phosphonates can be an alternative P source to microorganisms [14]. Although the utilization mostly occurs in phosphate-limited ocean regions, 2-AEP can be adopted by prokaryotes when DIP is sufficient as well [15, 16]. Catabolism of 2-AEP have been well elaborated in prokaryotes employing diverse pathways mediated by C-P lyase and C-P hydrolyses respectively [17, 18]. The C-P lyase pathway with broad substrate specificity is more commonly present under P deficiency condition to provide both P and sarcosine as reported in bacteria [19]. Substrate-inducible C-P hydrolase pathway can hydrolyze the phosphonates to release P, C, N, or energy sources [20]. Transportation of extracellular 2-AEP into the cells through the phosphonate transporter complex is believed to be the precondition to implement these pathways, referring to the case study that absence of transporters result in C-P hydrolyses pathway not functional in dinoflagellates [21].

Synthetic C-P compounds such as herbicides, insecticides, antibiotics [22] are widely used and the contamination of herbicides glyphosate (GLY) and glufosinate (GLU) have caused extensive concern [23, 24]. Whether the accumulation of these compounds in the coastal waters could be a toxin or a potential P eutrophic resource for marine organisms remains to be investigated. Efforts showed that several phytoplankton species are able to grow in the medium supplied with biogenic and synthesized phosphonate respectively, however leaving a big question mark about how it is utilized [25, 26]. In general, we have limited knowledge about the utilization of phosphonate and underlying mechanism in eukaryotic phytoplankton. Regarding that, the most pressing issues include, (1) the universality of the bioavailability of phosphonate compounds by eukaryotic phytoplankton, (2) the underlying mechanism of the assimilation or catabolism pathway.

Diatoms represent a major class of phytoplankton in the ocean, contributing ~20% of the primary production [27, 28]. *Phaeodactylum tricornutum* is a model diatom species extensively studied because of its biological characters, and it was reported being able to utilize synthetic phosphonate compound glyphosate [25, 29]. The completion of whole genome sequence of *P. tricornutum* lay the foundation to illustrate the metabolic mechanism at the cellular level [30]. In this study, we firstly investigated the bioavailability of biogenic phosphonate 2-AEP and synthetic phosphonates glyphosate (GLY), glufosinate (GLU) by *P. tricornutum*. Then we conducted comparative transcriptomic analysis to unveil the metabolic pathway regarding to the utilization of extracellular 2-AEP by *P. tricornutum*. Based on that result, we carried out further exploration of environmental meta-omics data analysis together with batch culture experiments to justify the proposed mechanism and discuss the implication in adaptive capacity.

## Methods

### Culture conditions and experimental setup

In this study, we conducted progressive experiments in three batches to address specific issues respectively (culture condition provided in Supplementary Table S1). Before the experiment, *P. tricornutum* seed culture was subjected to phosphorus starvation to deplete the intracellular P storage for 8-10 days while the DIP concentration in the medium was below the detection limit (~0.5 μM). The first batch culture was aimed to investigate bioavailability of different phosphonate compounds by *P. tricornutum*. The second batch culture was designated to explore the 2-AEP uptake efficiency of *P. tricornutum*. The third batch culture was set up to examine the cell response to different temperature while supplied with 2-AEP.

### Determination of physiological parameters (cell density, Fv/Fm, APA)

Cell density and Fv/Fm were measured daily while the alkaline phosphatase activity (APA) was determined every other day during the course of the experiments. The cell density was monitored using CytoFLEX flow cytometer (Beckman Coulter, Indianapolis, IN, USA) (detail provided in Supplementary). The photochemical efficiency of photosystem II Fv/Fm was measured daily using a FIRe Fluorometer System (Satlantic, Halifax, NS, Canada) as described earlier [31]. As a biomarker for P-stress, APA was determined by colorimetric method using p-Nitrophenyl Phosphate (pNPP) (Sigma-Aldrich, St. Louis, MO, USA) as the substrate according to the method described before [32].

### Determination of phosphorus concentration, cellular carbon and nitrogen contents

Cellular carbon and nitrogen contents were determined using a vario EL cube analyzer (Elementar Analysensysteme GmbH, Hanau, Germany) [33] (Supplementary). The carbon and nitrogen content were calculated as per cell and C/N ratios were obtained. DIP was measured daily to monitor the scarce of phosphate (Pi) in P-depleted and phosphonate groups using molybdenum blue method [34]. DOP measurement was carried out every other day during the course of the experiments. We established a high performance liquid chromatography (HPLC) based method for the determination of three phosphonates (GLY, GLU and 2-AEP) in salt matrix, which will be reported elsewhere (brief description provided in Supplementary).

### RNA-seq analysis and quantitative reverse transcription PCR (RT-qPCR)

For each sample, total RNA of 10^8^ cells was extracted as previously reported [35] and stored at −80 °C for subsequent cDNA synthesis and high-throughput sequencing. In total, 24 cDNA samples representing different cell growth status under different P nutrient condition (Supplementary Table S1) were subjected to RNA-seq sequencing (BGI, Wuhan, Hubei, China). After filtration, obtained clean reads were assembled and blast against the reference genome of *P. tricornutum* and reference gene respectively. The raw sequence reads have been submitted to the SRA at NCBI under BioProject ID is PRJNA764555. Differentially expressed genes (DEGs) were identified and mapped to Gene Ontology (GO) and KEGG annotation. Selected genes were subjected to RT-qPCR to verify transcriptional expression. Bioinformatic pipelines used in reads filtration, sequence annotation, identification of DEGs and methods of RT-qPCR were provided in Supplementary.

### Phylogenetic analyses and biogeographic distribution of selected genes

Selected genes representing key players in the proposed metabolism pathway were subjected to further analysis. To examine the prevalence and phylogenetic relationship of selected target genes among diatom and other eukaryotes, the deduced amino acid sequences from representative organisms were retrieved for subsequent phylogenetic tree reconstruction. We conducted further investigation of the active expression and biogeographic distribution of selected genes in the natural plankton assemblages through exploration of Ocean Gene Atlas (OGA) database [36]. (Method details provided in Supplementary).

### Transmission electron microscopy (TEM)

*P. tricornutum* grown with different phosphorus nutrient under different temperatures (12 °C and 25 °C) (Supplementary Table S1) were prepared for TEM imaging by using Tecnai G2 Spirit BioTwin Transmission Electron Microscope (FEI, Eindhoven, Netherlands) (Method details provided in Supplementary).

### Data statistical analysis

To determine whether the difference between different groups is statistically significant, we applied T-tests to calculate the P-value. Statistical analysis about RNA-seq data were described in Supplementary. Log2 transformed FPKM and RPKM were used to draw the heatmap and scatterplot, respectively. Besides, 1 was added to each FPKM and RPKM value before log2 transformation to facilitate calculation.

## Results

### Promoted cell growth and physiological responses in 2-AEP culture

In contrast to arrested growth in P− group, among four examined phosphonate culture conditions, we observed comparable cell growth in 2-AEP group (36 μM) and the phosphonate mixture group (36 μM) indicating that only 2-AEP can be utilized as the alternative P source by *P. tricornutum*, while GLY and GLU failed (Fig. 1a, Supplementary Fig. S1a). In comparison with the rapid growth in both DIP groups, cells exhibited moderate growth in cultures containing 36 μM 2-AEP, reached maximum cell density about half of that in 36 μM DIP group on day 3, then remained stable. In accordance with the cell growth, Fv/Fm representing the photosynthetic capacity exhibited the similar trend (Fig. 1b, Supplementary Fig. S1b). Significant increase (p<0.05) of Fv/Fm value was observed in DIP (36 μM) group (Fig.1b), and no difference identified between P-depleted group and the other groups (Supplementary Fig. S1b). In 2-AEP groups, Fv/Fm increased slightly, peaked on day 3 (close to the value of DIP 3.6 μM), then decreased to the same level of P− group (Fig. 1b). Starting from C/N ratio ~6.8, DIP (36 μM) group possessed the lowest value ~5.6, the highest value ~8.4 was observed in the P− group, and cultures of both 2-AEP groups and DIP (3.6 μM) group shared the median value ~7.5 (Fig. 1c).

**Fig. 1.**
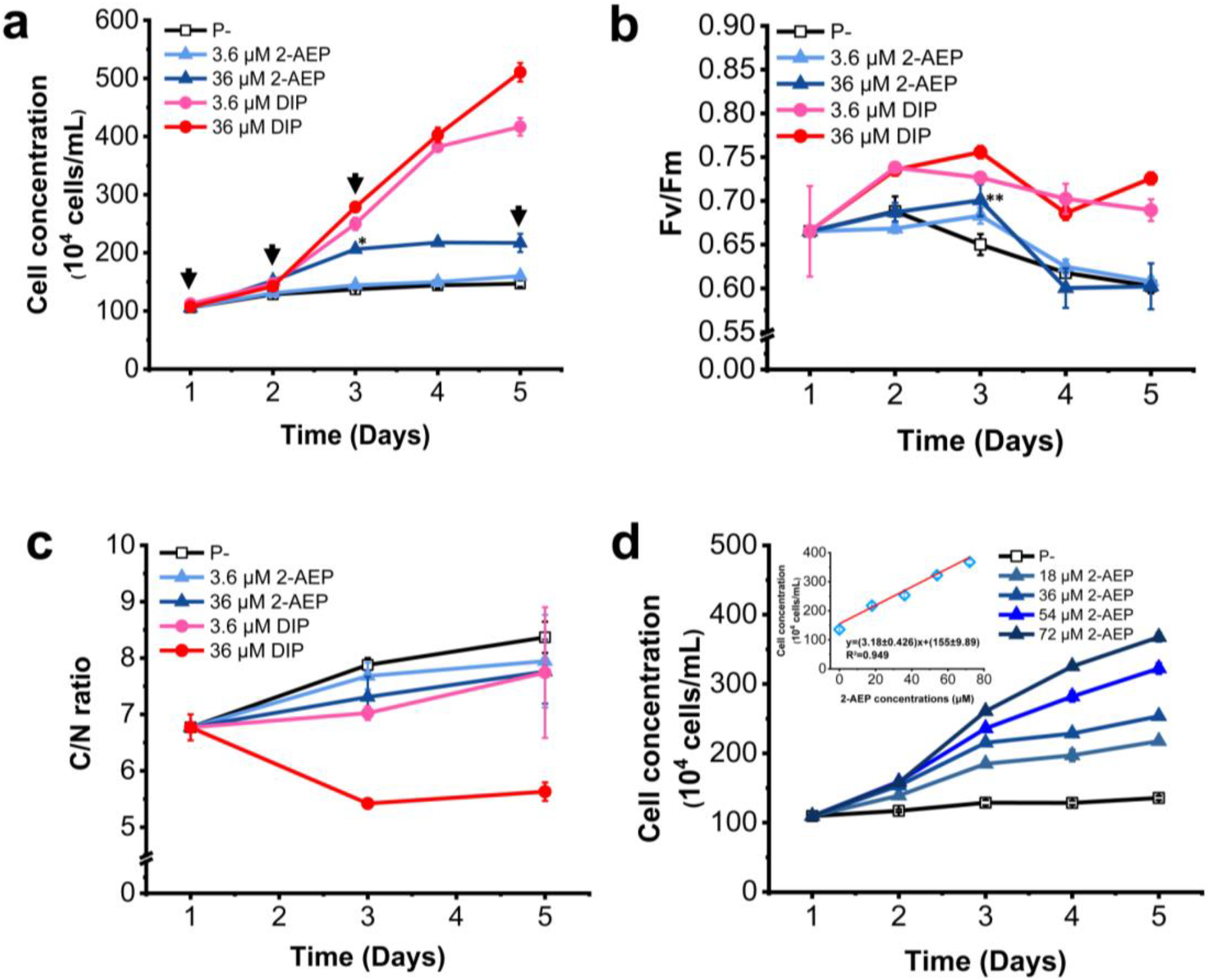
Physiological responses of *P. tricornutum* under different phosphorus conditions. (a) Growth curves, (b) Fv/Fm, (c) C/N ratio, (d) cell concentration grown under different 2-AEP concentration and correlation between 2-AEP concentration and *P. tricornutum* cell density on D5 shown in up-left. Data are averages of biological triplicates; error bars show ± S.D. of biological triplicates. P-value was calculated by T-test to represent significant difference between 36 μM 2-AEP group and P− group in (a) (b), * p=0, ** p=0.031.

### Stochastic uptake of 2-AEP by *P. tricornutum*

Briefly, physiological parameters showed that the utilization of 2-AEP is comparable to DIP (3.6 μM) by *P. tricornutum*, presented by similar Fv/Fm and C/N ratio. Meanwhile, elevated APA (Supplementary Fig. S1c) and arrested cell growth after day 3 indicates that utilization is incomplete given that 2-AEP was provided in higher concentration. Thus, we set up a gradient culture to further explore the utilization of 2-AEP (Supplementary Table S1). Results showed that stimulated cell growth is positively correlated with the increase of ambient 2-AEP concentration in a linear relationship (Fig. 1d). After 5 days, we observed highest cell concentration in 72 μM 2-AEP group, with about 4 times of that in the beginning, a stimulation comparable to that observed in 3.6 μM DIP (Fig. 1a, d). Moreover, concentrations of 2-AEP in the culture medium decreased both in 36 μM and 72 μM treatment which indicated that the 2-AEP were absorbed by *P. tricornutum* (Supplementary Fig. S1d).

### Overview of comparative transcriptomics and metabolic activity differences

After filtration, high-quality reads were assembled and mapped to the reference genome with the average mapping rate of 86.61% (Supplementary Table S2). A total of 11,293 genes was detected, including 11267 known genes and 26 unknown genes. Independent comparison within two 2-AEP groups revealed that most detected genes are shared by both (Supplementary Fig. S2a, b). Samples of D5 of both 2-AEP groups (3.6 and 36 μM) were subjected to further comparative analysis against P− and P+ groups showing 10561 genes shared among 4 groups (Fig. 2a).

**Fig. 2.**
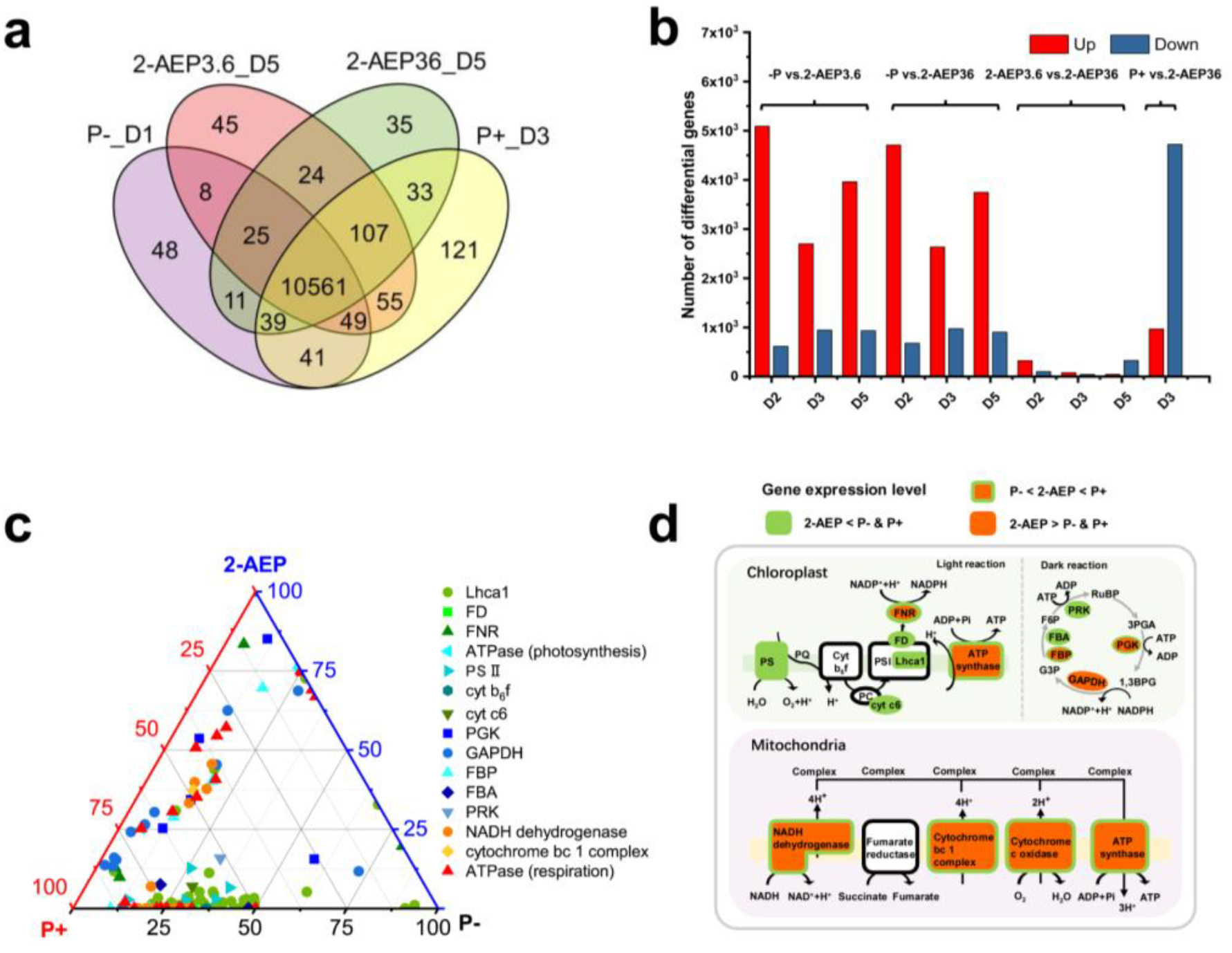
Comparative gene expression profiles of P. tricornutum under different phosphorus conditions. (a) Venn diagram showing shared genes among 2-AEP groups (3.6 μM and 36 μM, D5), P-repleted (P−, D1) group and P-replete (P+, D3) group. (b) Pairwise comparison of DEGs between different groups. (c) Ternary plots showing the DEGs of photosynthesis and respiration in P−, P+ and 2-AEP36_D5 group. DEGs include LHCA1 (LHCA1), PS II (psbO, psbU, psbQ, psbM, psb27, psbP), FD (petF), FNR (petH), cyt b6f (petC), cyt c6 (petJ), ATP synthase in photosynthesis (ATPF1A, ATPF1G), PGK (PGK), GAPDH (*GAPDH*), FBP (*FBP*), FBA (*FBA*), PRK (*PRK*), NADH dehydrogenase (*NDUFA5, NDUFAB1, NDUFA2, NDUFS5, NDUFB7, NDUFB9, NDUFV2, NDUFA12, NDUFB10, NDUFS6, NDUFA9*), cytochrome bc1 complex (*QCR7*), ATP synthase in respiration (*ATPeV1H, PMA1, ATPF1A, ATP5O, ATP5D, ATP5A1, ATP5C1, ATP5E, ATP6C, ATPF1G, ATP6D, ATP6L, ATP6B, ATP6A, ATP6M, ATP6S14*). Each type of points corresponds to DEGs of components involved in photosynthesis and respiration. Its position represents its expression level with respect to each group (P−, P+,2-AEP36_D5). (d) Schematic summary of metabolic activity differences in photosynthesis and respiration in *P. tricornutum* under different phosphorus conditions. Specific information of DEGs in different groups refer to Supplementary Table S3.

Through pairwise comparison among different groups (Fig. 2b, Supplementary Table S3), we identified significant higher number of DEGs between P− vs. 2-AEP (3.6 and 36 μM) and P+ vs. 2-AEP (36 μM) group, but limited DEGs identified in 2-AEP groups comparison (3.6 μM vs. 36 μM). Most of the DEGs were up-regulated in 2-AEP cultures compared with P− group, while most DEGs identified as down-regulated compared with P+ group (Fig. 2b). Gene Ontology (GO) categorization showed that detected DEGs (P− vs. 2-AEP36_D5) were significant enriched in cellular process, metabolic process, membrane, binding and catalytic activity (Supplementary Fig. S2c). Regarding KEGG pathways mapping, most DEGs were enriched in the pathway related to metabolism, including lipid metabolism, energy metabolism and global and overview maps (Supplementary Fig. S2d).

Generally, in-depth analyses of cellular metabolic activity among these groups are consistent with observed physiological responses, showing moderate gene transcriptional level of photosynthesis and respiration in 2-AEP groups compared with P− and P+ groups (Fig. 2c, d, Supplementary Table S4).

The light reaction of photosynthesis is composed of photosystem (PS II and PS I), electron transport chain (ETC) and generation of ATP [37]. Transcriptional expression of genes involved in PS II (*psbO, psbU, psbP, psbQ, psbM, psb27*), PS I (*LHCA1*) and ETC (FD (*petF*)) were repressed in 2-AEP groups against P− and P+ groups (Fig. 2c, d, Supplementary Table S4). Gene expression levels of *FNR* (*petH*) and *ATPase* (*ATPF1A, ATPF1G*) of 2-AEP groups were situated between P− and P+ group (Fig. 2c, d, Supplementary Table S4). In the dark reaction where carbon fixation occurs, *PGK* and *GAPDH* involved with reduction were up-regulated compared to P− group. Whilst, the genes *FBA* and *PRK* involved with RuBP regeneration were down-regulated compared to both P− and P+ group (Fig. 2c, d, Supplementary Table S4). Gene transcripts associated with oxidative phosphorylation of mitochondria (NADH dehydrogenase, cytochrome bc 1 complex, cytochrome c oxidase, ATP synthase) exhibited similar expression pattern, that lies between P− and P+ group (Fig. 2c, d, Supplementary Table S4).

Furthermore, comparable up-regulation of P-stress marker genes were identified in 2-AEP groups and P− group in comparison with P+ group (Supplementary Table S5). In accordance with elevated APA in the culture, AP genes *PhoA* (Phatr3_J47869) and *PhoD* (Phatr3_J45959) was upregulated 4.45 and 2.49 Log2FC respectively (P+ vs. P−), 6.39 and 2.57 (P+ vs. 2-AEP36_D3), sharing similar value to the reported value 6.84 and 2.09 Log2FC [38]. Phosphate transporter genes *NPT* (Phatr3_J40433, Phatr3_J47667) exhibited1.51 and 2.95 Log2FC (P+ vs. P−), 1.98 and 2.92 (P+ vs. 2-AEP36_D3), similar to previous study as well [38]. Above results indicated that cells grown in 2-AEP group exhibited a metabolic reconfiguration despite under P-stress as P-depleted group. Guided by that, we made further efforts to decipher the underlying molecular mechanism involved in 2-AEP utilization by comparing 2-AEP groups against P-depleted group.

### Uptake of 2-AEP through clathrin-mediated endocytosis

Barely detectable DIP in 2-AEP culture suggested 2-AEP was transported into the cells to be utilized. However, genes coding phosphonate transporters PhnCDE, PhnSTU, AepXVW, AepP and AepSTU reported previously [16] were not identified in all examined groups (absence in the transcriptome). Instead, through integrated analysis, we deduced that 2-AEP was transported into cells encapsuled by the clathrin-coated vesicles.

Clathrin is a self-assembling protein that can form a cage-like lattices at the plasma membrane to perform vesicular uptake [39]. Clathrin-mediated endocytosis is a classical manner adopted by eukaryotes to transport cargo molecular into cells, characterized by a well-orchestrated process including coating (activation, vesicle formation) and uncoating. We identified the up-regulation of a whole train of genes involved with this process in 2-AEP groups compared with P− group (Fig. 3a, b, c, Supplementary Fig. S3a).

**Fig. 3.**
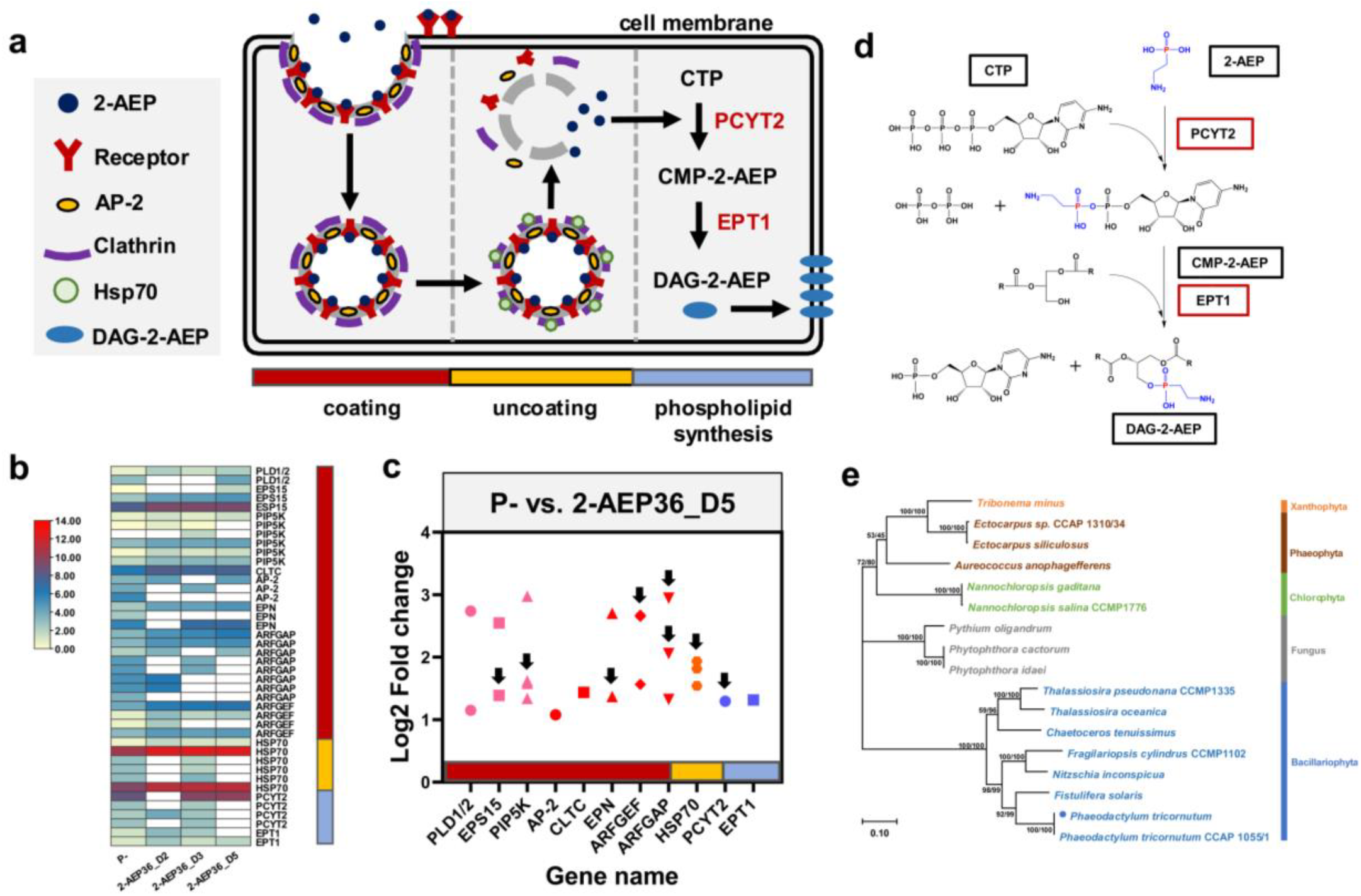
(a) Schematic summary of metabolic pathway of 2-AEP utilization by *P. tricornutum*. Coating process, 2-AEP encapsuled by the clathrin-coated vesicles, receptor genes including *EPS15, PLD1/2, PIP5K*. Uncoating process, 2-AEP released from the vesicles driven by HSP70 binding. Phospholipid synthesis, incorporation of 2-AEP into cell membrane catalyzed by two enzymes, ethanolamine-phosphate cytidylyltransferase (PCYT2) and Ethanolaminephosphotransferase (EPT1). CMP-2-aminoethylphosphonate (CMP-2-AEP), Diacylglyceryl-2-aminoethylphosphonate (DAG-2-AEP). (b) Expression level of multi copy genes in time series samples of 2-AEP36, (P−_D1 represent the initial cell condition), color bar represents the normalized gene expression level. *ARFGAP* (*SMAP, AERGAP1*), *ARFGEF* (*ARFGEF, GBF1*), *HSP70* (*HSPA1s*). (c) Fold changes of selected genes involved in endocytosis and phospholipid synthesis in group P− vs. 2-AEP36_D5, gene *EPS15, CLTC, HSPA1s, PCYT2* and *EPT1* were verified by qPCR (Supplementary Fig. S3a), *ARFGAP* (*SMAP*), *HSP70* (*HSPA1s*). Arrows represent genes up-regulated in P+ vs. 2-AEP group as well. (d) Phospholipids biosynthesis pathway show the incorporation of 2-AEP. Black boxes represent compounds, red boxes represent enzymes. (e) Phylogenetic tree inferred from amino acid sequences of clathrin of *P. tricornutum* and other eukaryotic organisms (Sequence IDs listed in Supplementary Table S7). Tree topology shown as generated from ML. Support nodes obtained from ML/NJ are shown as percentage. The blue dot represents the species used in this study.

Firstly, membrane coating process, including activation and vesicle formation is accomplished through a complex protein interaction network. *EPS15, PLD, PIP5K* are receptor genes coding for the protein responsible for endocytosis initiation, cargo recruitment and signal transduction respectively [40–42]. We identified 1.1~3.5 log2FC of gene *PLD1/2, EPS15* and *PIP5K* in both 2-AEP groups (D2, 3, 5) compared with that in P-depleted group (Fig. 3b, c, Supplementary Table S4). To be concise and specific, the expression level of these multi copy genes in time series samples of 2-AEP36 group exhibited similar pattern, most genes reached a highest transcriptional level on D5 (Fig. 3b, c). After the activation, *AP-2, Clathrin (CLTC)* and *EPN* genes were activated to form endocytic cups, upregulated within the range (1.0~2.7 Log2FC) in all the 2-AEP samples compared with that in P− group (Fig. 3a, c, Supplementary Table S4) [43–45]. Different from those genes, *ARFGEF* and *ARFGAP* (*SMAP*) genes playing important role in the membrane trafficking and actin remodeling [46, 47] were upregulated mostly on D2, and *ARFGEF* maintained higher expression till D5 with 1.6~2.7 Log2FC (Fig. 3b, c, Supplementary Table S4).

Regarding the uncoating process, heat-shock protein HSP70 is responsible for binding to clathrin and driving the dissociation of coated vesicles [48]. Results showed that *HSP70* (*HSPA1s*) gene expression was significantly up-regulated (1.5~2.7 Log2FC) in both 2-AEP groups (D2, 3, 5) than P− (Fig. 3a, b, c, Supplementary Table S4). The up-regulation of selected genes *EPS15* (Phatr3_J42442), *CLTC* (Phatr3_EG01984), *HSPA1s* (Phatr3_J54019) was further verified by qRT-PCR in the case of P− vs. 2-AEP36_D5 (Supplementary Fig. S3a, Supplementary Table S6). Besides that, we also identified the upregulation of selected genes (indicated by arrows in Fig. 3c) while compared with P+ group (Supplementary Fig. S3b, Supplementary Table S4).

### Recruitment of 2-AEP into phospholipids

After endocytosed into *P. tricornutum* cells, we proposed that the 2-AEP was catalyzed to be incorporated into the phospholipids (Fig. 3a, d). According to KEGG pathway (ko00440), 2-AEP firstly react with cytidine triphosphate (CTP) through the catalysis of ethanolamine-phosphate cytidylyltransferase (PCYT2) to produce cytidine monophosphate-2-aminoethylphosphonate (CMP-2-AEP) and diphosphate. Subsequently, CMP-2-AEP and 1,2-Diacyl-sn-glycerol were catalyzed by ethanolaminephospho-transferase (EPT1) to form diacylglyceryl-2-AEP (DAG-2-AEP), a component of membrane phospholipids (Fig. 3d). PCYT2 and EPT1, two key enzymes catalyzing the biosynthesis of DAG-2-AEP were significantly up-regulated of 2-AEP36 group 1.3~4.5 Log2FC compared with P− group and P+ group on D5 (Fig. 3b, c, Supplementary Fig. 3b, Supplementary Table S4). The differential expression of gene *PCYT2* (Phatr3_J40163) and *EPT1* (Phatr3_J37086) in P− vs. 2-AEP36_D5 group was verified by qRT-PCR as well (Supplementary Fig. S3a).

### Prevalence and transcript abundance of selected genes in environmental samples

Genes *CLTC, AP-2, PCYT2, EPT1*, representing endocytosis and incorporation process of 2-AEP utilization by diatom, were selected for further analysis. We identified the widespread of selected genes in reported diatom genome, including *P. tricornutum, Fistulifera solaris, Nitzschia inconspicua, Fragilariopsis cylindrus, Chaetoceros tenuissimus, Thalassiosira pseudonana* and *Thalassiosira oceanica*. Further phylogenetic analysis of deduced amino acid sequences derived from diatom and other representative eukaryotic organisms showed that selected genes of diatom form an independent branch in the phylogenetic tree (Fig. 3e, Supplementary Fig, S4).

Promoted by this result, we conducted a further exploration of biogeographic distribution of these selected genes involved with above-described mechanism in the environmental meta-omics dataset, Ocean Gene Atlas [14, 49]. We found a ubiquitous distribution of these genes in diatom assemblage worldwide, in both surface and DCM water layers (Fig. 4a). In general, selected genes actively expressed globally, showing similar biogeographic distribution pattern, to be specific, relative lower abundances detected in DCM layer compared to surface layer (Fig. 4a, Supplementary Table S8). Plotting abundances of selected gene transcripts from surface samples against in-situ temperature and phosphorus concentration showed that significantly enriched distribution (85% of total stations) of selected genes in waters with mild temperature ranging from 15 ~ 30 °C and low phosphate concentration (0 ~ 0.5 μM) (Fig. 4b, Supplementary Table S8). Furthermore, we observed that elevated abundances of individual samples move towards colder water (<10 °C) with higher phosphate concentration (> 1μM). Samples with highest transcript abundance of all selected gene transcripts were collected at the Southern Ocean (62.14°S, 49.3273°W) with highest phosphate concentration (~ 2 μM) and lowest temperature (~ 0°C) (Fig. 4b, Supplementary Table S8). Through searching the Southern Ocean (SO) diatom assemblage transcriptome database [50], we also identified abundant transcript expression of four selected genes (Supplementary Table S9) in diatom species shown in Fig. 3e.

**Fig. 4.**
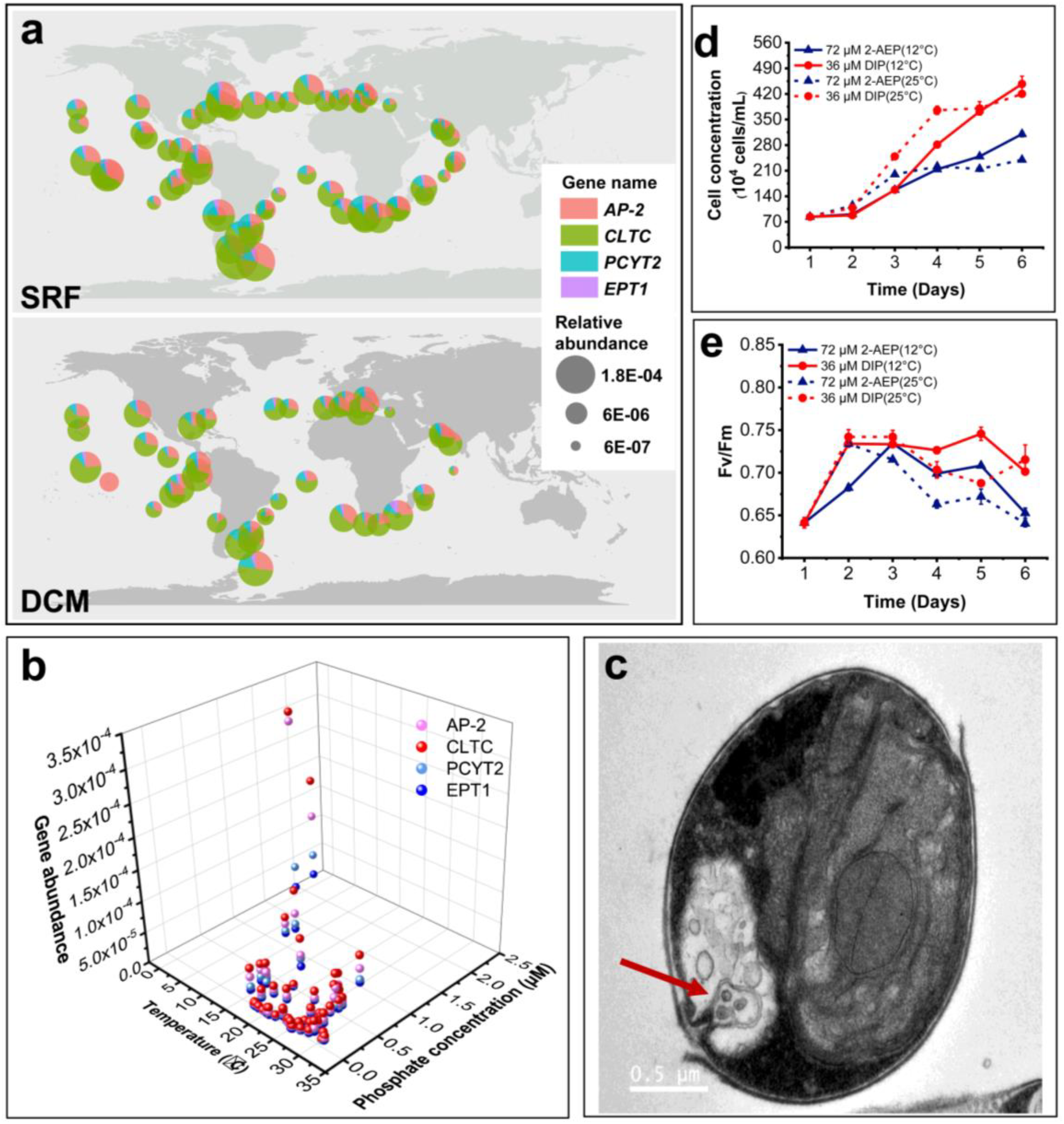
(a) Biogeographic distribution of gene *AP-2, CLTC, PCYT2* and *EPT1* of diatom in the ocean include the layer of surface (SRF) and the deep chlorophyll maximum (DCM). (b) Distribution of selected genes involved in 2-AEP utilization with respect to environment parameters (temperature and phosphate concentration) in surface waters (5 m depth). (c) Transmission electron microscopy (TEM) micrographs presenting the cell contain membrane bound vesicle in 2-AEP group (72 μM, 12°C). Cell growth (d) and Fv/Fm (e) of *P. tricornutum* under different phosphorus and temperature conditions.

### Vesicle image and cell growth under lower temperature in 2-AEP culture

Inspired by above results, we set up the third batch culture (Supplementary Table S1) to inspect the proposed mechanism and potential adaptive capacity. A cell with formed intracellular membrane-bound vesicle was clearly imaged by TEM (Fig. 4c) in 2-AEP culture grown at 12 °C (a potential counterpart at 25 °C), but fail to be spotted in DIP groups (Supplementary Fig. S5a, b, c).

Results showed that cell growth was promoted significantly in all groups, yet can be divided into two types. In the beginning, cell growth showed a day lag in both 2-AEP and DIP cultures grown at 12°C compared with instant increasing cell density in cultures at 25 °C (Fig. 4d). Despite that Fv/Fm in 2-AEP culture at 12 °C increased slower compared with other groups, it reached the maximum value on day 3 and maintained higher than that of 25 ° C 2-AEP culture (Fig. 4e). Elevated photosynthesis capacity is consistent with the higher cell density observed in 2-AEP (12 °C) than that in 2-AEP (25 °C). More than that, in 2-AEP cultures, both APA and C/N ratio possessed lower value at 12 °C against 25 °C, indicating that cells were less phosphorus stressed (Supplementary Fig. S5d, e). Accordingly, 2-AEP concentration decreased dramatically on D3 and remained unchanged till D5 at 12 °C (Supplementary Fig. S6f). Gene *EPS15, CLTC* and *HPS70* that involved in endocytosis were all upregulated on D3 at 12 ° C compared with 25 ° C (Supplementary Fig. S6g). Taken together, physiological parameters and gene expression suggested cells play more actively under lower temperature when supplied with 2-AEP.

## Discussion

### Cargo transportation through clathrin-mediated endocytosis

Clathrin-mediated endocytosis transports different kinds of extracellular molecules into the cells through vesicular trafficking [51]. This process, requiring different components work together to drive the formation and dissociation of vesicles, has been elucidated in many eukaryotic model organisms sharing similar modules. Yet up to now, there is limited reports describing this mechanism in phytoplankton, to which uptake nutrient from environment is crucial for their growth and primary production in the ocean. Annenkov proposed that the diatom may capture silicon by endocytosis hypothetically, yet lacking experimental evidence [52]. Turnšek’s research hypothesized that the iron was transported by phytotransferrin which are endocytic into the diatom cells [53]. Shemi’s research identified that phosphorus starvation enhanced the formation of membrane vesicles in the cytoplasm of *Emiliania huxleyi* [54]. Besides, in dinoflagellate *Prorocentrum donghaiense* and green algae *Chlamydomonas reinhardtii*, this process has been reported to be a route for both nutrient and harmful substances to enter the algal cells, respectively [55, 56]. Though, the integrity pathway of this mechanism and dynamic expression under different nutrient condition was not elaborated before.

Clathrin-mediated endocytosis is considered to be a random process [57]. The fundamental principle of cargo recruitment is that the protein components of clathrin coat bind to specific binding sites of different transmembrane cargo molecules to recruit them to the region in cytomembrane that will form the vesicle, then the enriched cargos in the forming vesicles being selectively endocytosed [58]. If the cargo molecules have more opportunity to combined with the adaptors namely protein components of clathrin coat, the likelihood of endocytosis initiation would be increased [57]. In this study, this hypothesis is supported by the gradient culture experiment. Attributed to higher concentration, the opportunity to form vesicle coat increased as well, showing that stimulated cell growth in accordance with the increase of 2-AEP concentration in ambient conditions in a linear relationship (Fig. 1d).

The worldwide distribution and enriched expression of endocytosis key genes in low phosphate regions suggest that this mechanism might be a prevalent strategy employed by diatom in pelagic environment. Characterized by randomness, it is reasonable to deduce that diverse forms of nutrients could be adopted through the vesicle mechanism as well. Diatoms are identified as autotrophic or mixotrophic, but the underpinning mechanism of heterotrophic remain unclear [59]. This study indicate that diatom might be able to grow in heterotrophic mode with the aid of clathrin-mediated endocytosis hypothetically.

There are different ways of vesicle formation in endocytosis, and clathrin-coated vesicle is reported to have the larger average size about 100-200 nm in diameter, that is consistent with our observation (~200 nm, Fig. 4c) [60, 61]. Ultrastructure of the imaged vesicle (Fig. 4c) was slightly different from the typical clathrin-coated vesicle featured by surrounding coated pellets in deeper color. The possible reason might be that, we only identify the Clathrin heavy chain coding gene in *P. tricornutum*, with the absence of light chain coding gene that fail to form the denser cage structure. Nevertheless, the comparable vesicle size captured in 2-AEP culture together with the transcriptional regulation provided the evidences about the formation of clathrin-mediated endocytosis.

### Incorporation of 2-AEP in membrane phospholipid and physiological implication

Synthesis of phospholipids and nucleic acids are the major phosphorus requirement in phytoplankton cell growth [62]. Phospholipids are main constituents of cell membranes and glycerophospholipids accounts for the vast majority of phospholipids [63], and glycerophospholipids are especially important for being structural and functional components of cell membrane [64]. Study shows that 2-AEP firstly reacts with CTP to form CMP-2-AEP, which then is transferred to diglyceride to form the glycerophospholipid [5]. The process of DAG-2-AEP synthesis has been described in animal tissues [65, 66]. Though there is no report about the presence of DAG-2-AEP in algae cells, we are able to depict the biosynthesis pathway based on the transcriptomic data obtained in this study. The up-regulation of two enzymes which catalyze the incorporation of DAG-2-AEP from 2-AEP lead to a hypothesis, the *P. tricornutum* use 2-AEP to synthesis cell membrane lipids to maintain the cell structure and morphology, which enable the cells to reallocate the phosphorus distribution among cellular metabolisms under P-stress condition. Ostrowski’s research identified elevated 2-AEP amounts and redistribution of lipids in the fusion site during *Tetrahymena* mating provide evidence to support our claims [67]. Besides that, our results showed that cellular C/N ratio of *P. tricornutum* were deeply affected by P nutrient condition, the lowest value 5.6 (P+) in consistent with the value 5.64 reported in *P. tricornutum* under P sufficient condition [68]. Median C/N ratio of 2-AEP group compared with P-depleted group also indicated that cells were partially relieved from P stress and managed to adjust the cellular resource allocation.

We observed impeded photosynthesis but elevated respiration in 2-AEP group, suggesting that higher demanded in energy (Fig.1b, Fig. 2c, d) which may be consumed by the endocytosis and membrane incorporation processes. All these results showed that 2-AEP is an alternative phosphorus source instead of preferred phosphorus source to *P. tricornutum*.

In the light of many previous studies, bacteria and cyanobacteria are able to utilize 2-AEP by cleaving the C-P bond of 2-AEP to form phosphate [15, 16]. In contrast, the 2-AEP utilization mechanism of eukaryotic *P. tricornutum* unveiled in this study is disparate. 2-AEP is chemically stable and its dissociation energy is much higher than that of other organic phosphorus [69]. To cleave the C-P bond of 2-AEP, *P. tricornutum* might consume much more energy than incorporated them into the phopsholipids of cell membrane. Regarding the energy consumption, the mechanism described in this study would be a cost-effective approach for *P. tricornutum* to survive under phosphate limitation condition.

Furthermore, at the community level, higher relative abundances of selected genes identified in cold waters and enriched distribution in low P waters, leading to a speculation that the utilization of 2-AEP could be adopted as a survival strategy and defense mechanism for diatoms grown under stress condition caused by nutrient deficiency or low temperature, which is then proved by our further culture experiment. At present, P deficiency in the ocean was prevalent, and the DOP concentrations in pelagic ocean are higher than DIP concentrations especially in the surface water [70]. Our results showed relative higher expression of selected genes in surface samples is consistent with this pattern. Under these circumstances, the phosphonate bioavailability and utilization exemplified by diatom should be further examined to evaluate the contribution to biomass, primary production and the biogeochemical cycle of dissolved and particulate P pools in the ocean.

## Conclusion

Firstly, this study showed that biogenic phosphonate 2-AEP can promote the growth of *P. tricornutum* under P deficiency. Through comparative omics analysis, we elaborated an unconventional mechanism adopted by *P. tricornutum* to utilize extracellular 2-AEP. We proposed that 2-AEP is transported into the cells through clathrin-mediated endocytosis and then incorporated into the membrane phospholipids. Active expression of selected genes representing this deductive mechanism are ubiquitous in the ocean and enriched in the regions with moderate temperature and low phosphate concentration according to the analysis of environmental meta-omics dataset. Taken together with batch culture experiments, our findings indicated that the 2-AEP can be an alternative phosphorus source for *P. tricornutum* and the utilization of 2-AEP might play important role in facilitating adaptive capacity. The ecological implications of this proposed mechanism of diatom requires further rigorous experimental verification. With the combination of isotopic-label tracer technique and glycerophospholipid HPLC, we are expecting to inspect the proposed mechanism in vivo and in-situ. Overall, this study unveils a new chapter and provides the insight for future endeavor in exploring the utilization of phosphonate by eukaryotic phytoplankton.

## Acknowledgements

This research was supported by the National Natural Science Foundation of China grants # 41776106 (J.M.), # 41706116 (X.L.), and Natural Science Foundation of Fujian Province of China # 2020J06008 for Distinguished Young Scholars (J.M.). We are grateful to Senjie Lin and all members of Marine EcoGenomics Laboratory (Xiamen University) for their assistance in this study. We thank Xin Lin, Zichao Deng and Yawei Luo (Xiamen university) for the assistance in data analysis. We thank Junhui Chen (Xiamen university) for the instruction in carbon and nitrogen determination. We thank Lumin Yao, Caimin Wu, Qingbing Zheng (Xiamen university) for the assistance with TEM imaging. We thank Bin Geng (BGI Genomics Co., Ltd.) for assistance with RNA sequencing.

## Author contributions

Huilin Shu, Xin Lin and Jian Ma conceived this study. Huilin Shu, Jian Ma, Yanchun You and Ling Li conducted the lab work. Data analyses and manuscript were done by Huilin Shu, and Xin Lin, with input from Jian Ma, Hongwei Wang and Jingtian Wang. All authors edited the final version before they approved submission.

