## Supplementary materials and methods for "Comparative transcriptomics unveil distinctive metabolic pathway of phosphonate utilization by diatom *Phaeodactylum tricornutum*"

**Determination of cell density and APA**

1 mL cell suspension sampled in centrifuge tube was subjected to a total of 10,000 counting events for each measurement. The evaluation of cell density was determined by gating areas in the Chlorophyll A vs SSC-A dot plot in which all cells appear.

Another 1 mL sample was collected to determine APA. 50 μL PNPP was added to each sample. After 2 hours reaction, the absorbance of supernatant was measured at 405 nm. Different concentrations of PNP were used to making calibration curve and the APA was calculated as PNP concentration per cell per hour.

**Determination of phosphorus concentration, cellular carbon and nitrogen contents**

To determine 2-AEP concentration, the medium filtrate was firstly prepared and diluted to a proper salinity and concentration, then a derivatizing regent 9-fluorenylmethyl chloroformate (FMOC-Cl) at concentration of 6.0 mmol/L was used to react with the phosphonates in the diluted filtrate. After 30 mins of reaction, the derivatives were determined by liquid chromatography with fluorescent detection (excitation wavelength at 265 nm and emission wavelength at 315 nm).

Cellular carbon and nitrogen contents were determined using a vario EL cube analyzer (Elementar Analysensysteme GmbH, Hanau, Germany) [1]. 5 mL cell culture were collected every other day and filtered onto 25 mm GF/F glass fiber filters which had been precombusted at 450 °C for 4 h. GF/F filtrates were preserved at -20 °C. Before determinations, frozen filters were dried at 60 °C for 8 h. After that, 500 μL 1% HCl were dripped onto the filters and the filters were dried again at 60 °C for 12 h. The carbon and nitrogen content were calculated as per cell and C/N ratios were obtained.

**RNA isolation, RNA-seq and RNA-seq analysis**

10^8^ cells were collected by centrifugation at 6000 rpm for 10 min at 4 °C, then resuspended in 1 mL TRIzol (Invitrogen, Carlsbad, CA, USA) and stored at -80 °C before RNA extraction. For RNA isolation, a Direct-zol RNA Miniprep Kits (Zymo research, Irvine, CA, USA) was used and the extracted total RNA was dissolved in RNase-free water and stored at -80 °C for subsequent high-throughput sequencing. RNA-seq data analysis were conducted by BGI (Wuhan, Hubei, China).

6.37 Gb reads were obtained per sample after trimming the adaptors as well as removing low-quality sequences and unknown reads with extremely high N bases. The reference genome of *P. tricornutum* was downloaded from ftp://ftp.ensemblgenomes.org/pub/protists/release38/fasta/phaeodactylum_tricornutum/dna//Phaeodactylum_tricornutum.ASM15095v2.dna.toplevel.fa.gz. The clean reads were assembled and blast with the reference genome using HISAT2 (v2.0.4) [2] and reference gene using Bowtie2 (v2.2.5) [3] respectively.

The gene expression level was calculated using RSEM (v1.2.12) [4] and normalized to FPKM (Fragments per kilo base of transcript per million mapped reads) value. Genes with Log2FC (fold change) ≥1 and p-value ≤ 0.05 (adjusted p-value) were defined as significantly differentially expressed genes (DEGs) [5]. DEGs were subsequently mapped to Gene Ontology (GO) terms to determine the gene ontology and classified into different biological pathways according to the KEGG annotation by using Phyper (https://en.wikipedia.org/wiki/Hypergeometric_distribution) based on Hypergeometric test. The significant levels of terms and pathways were corrected by Q value with a rigorous threshold (Q value ≤ 0.05) by Bonferroni [6].

**Quantitative reverse transcription PCR (RT-qPCR)**

Specific primers (Supplementary Table S2) were designed based on the unigene sequence with highest fold-change obtained in DEGs profile. We used ribosomal protein coding gene *RPS*（Phatr3_J10847）as reference gene to normalize the expression of the target genes as reported [7]. RT-qPCR was performed on iCycle iQ Real-Time PCR Detection System using Bio-Rad iQ SYBR Green Supermix Kit (Bio-Rad Laboratories, Hercules, United States). The fold change of selected genes was determined by 2^(−ΔΔCt)^ [8].

**Phylogenetic analyses and biogeographic distribution of selected genes**

Deduced amino acid sequences from representative organisms were retrieved from NCBI and aligned using ClustalW on the MEGA X platform [9]. Phylogenetic tree reconstruction was performed using Maximum likelihood and Neighbor-joining method based on Jones-Taylor-Thornton (JTT) matrix model with 1000 bootstraps in MEGA X [9].

Nucleotide sequences of selected genes *AP-2* (Phatr3_J18142), *CLTC* (Phatr3_EG01984), *PCYT2* (Phatr3_J40163) and *EPT1* (Phatr3_J33864) were used as query to BLASTX against the Ocean Gene Atlas (OGA) database [10] with the cut off e-value 1E-10. Genes annotated as diatom were extracted and subject to further analysis of the global distribution pattern. Abundances were computed as RPKM (reads per kilobase covered per million of mapped reads) and the distribution pie chart were calculated by using ‘percentage of total abundance per gene’. The biogeographic distribution of selected genes in diatoms was plotted in R (v.4.1.1) using scatterpie and ggplot2 [11, 12]. Besides, the abundances of selected genes and expression of surface samples (5 m depth) were further analyzed in the context of environmental parameters through three dimensional scatter plot.

**Transmission electron microscopy (TEM)**

Cells were collected by centrifugation and fixed in 2.5% glutaraldehyde for 12h. Then, the cells were pre-embedded in 2% agarose gel. Subsequently, samples were dehydrated, infiltrated, embedded and sectioned by Leica UC7 Ultramicrotome (Leica Microsystems, Wetzlar, Germany) and observed by using a Tecnai G2 Spirit BioTwin Transmission Electron Microscope (FEI, Eindhoven, Netherlands).
