## Supplementary figures and tables for "Comparative transcriptomics unveil distinctive metabolic pathway of phosphonate utilization by diatom *Phaeodactylum tricornutum*"

### Slide 1
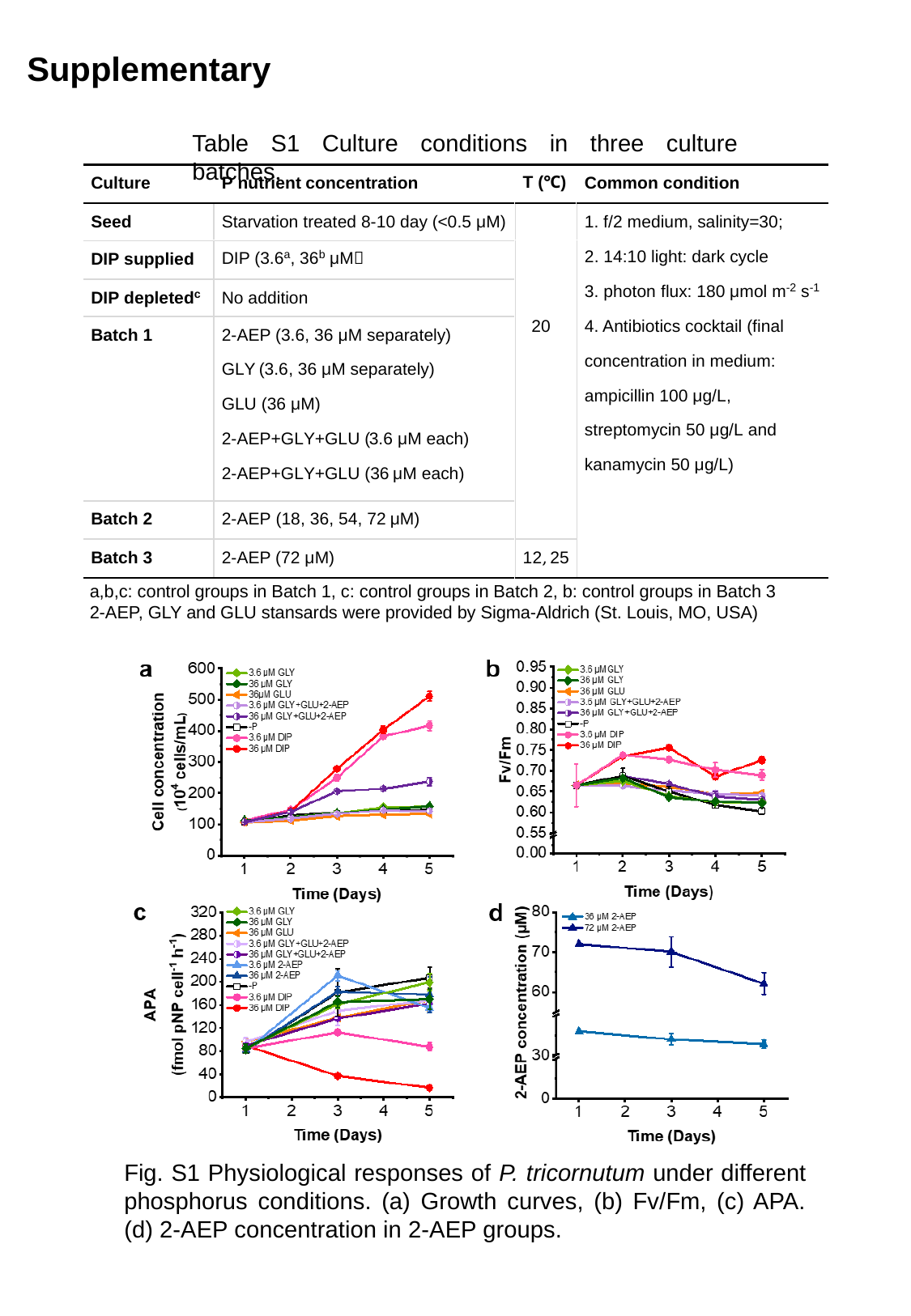

Supplementary
Table S1 Culture conditions in three culture batches.
a,b,c: control groups in Batch 1, c: control groups in Batch 2, b: control groups in Batch 3
2-AEP, GLY and GLU stansards were provided by Sigma-Aldrich (St. Louis, MO, USA)
Fig. S1 Physiological responses of P. tricornutum under different phosphorus conditions. (a) Growth curves, (b) Fv/Fm, (c) APA. (d) 2-AEP concentration in 2-AEP groups.

### Slide 2
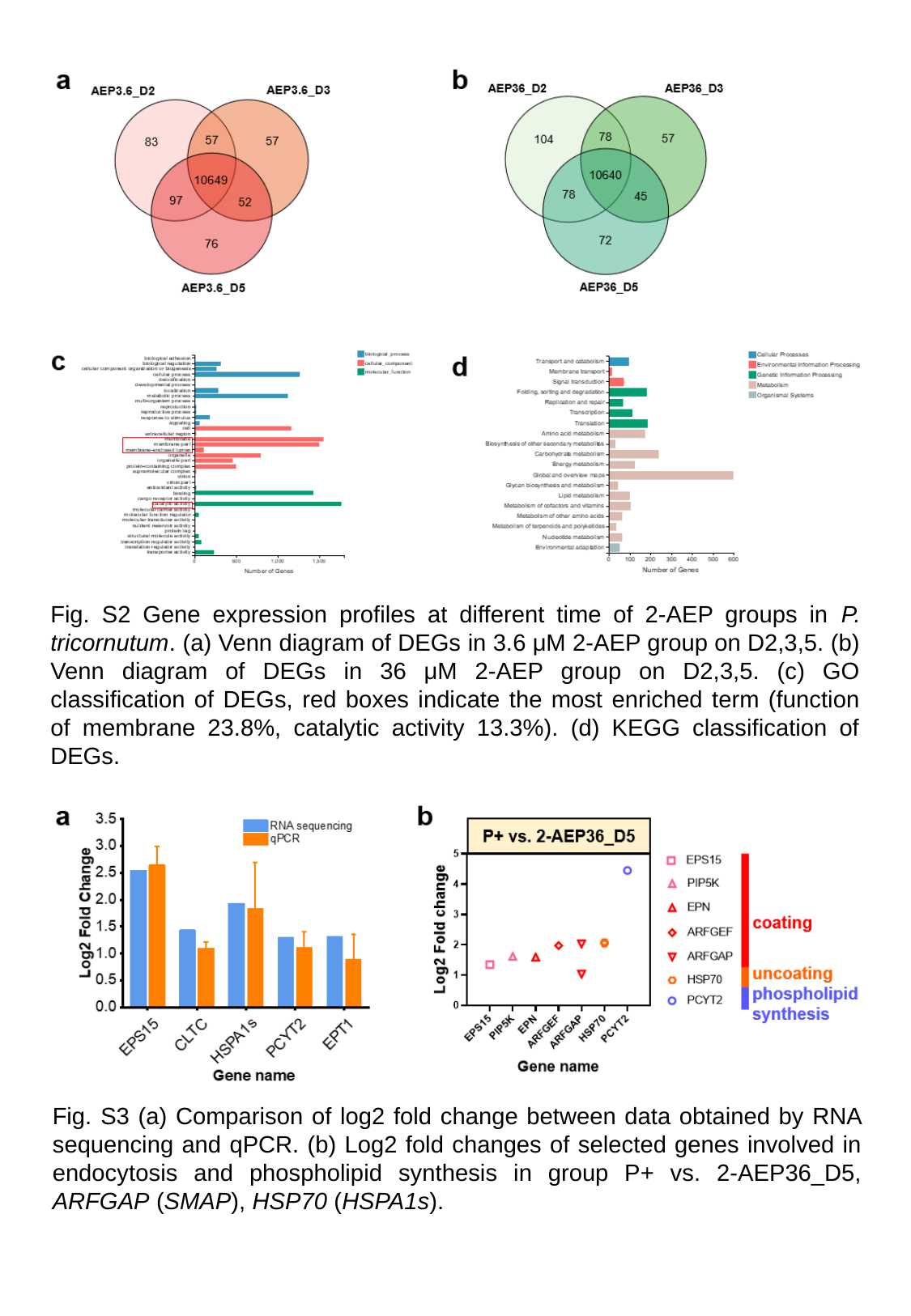

Fig. S2 Gene expression profiles at different time of 2-AEP groups in P. tricornutum. (a) Venn diagram of DEGs in 3.6 μM 2-AEP group on D2,3,5. (b) Venn diagram of DEGs in 36 μM 2-AEP group on D2,3,5. (c) GO classification of DEGs, red boxes indicate the most enriched term (function of membrane 23.8%, catalytic activity 13.3%). (d) KEGG classification of DEGs.
Fig. S3 (a) Comparison of log2 fold change between data obtained by RNA sequencing and qPCR. (b) Log2 fold changes of selected genes involved in endocytosis and phospholipid synthesis in group P+ vs. 2-AEP36_D5, ARFGAP (SMAP), HSP70 (HSPA1s).

### Slide 3
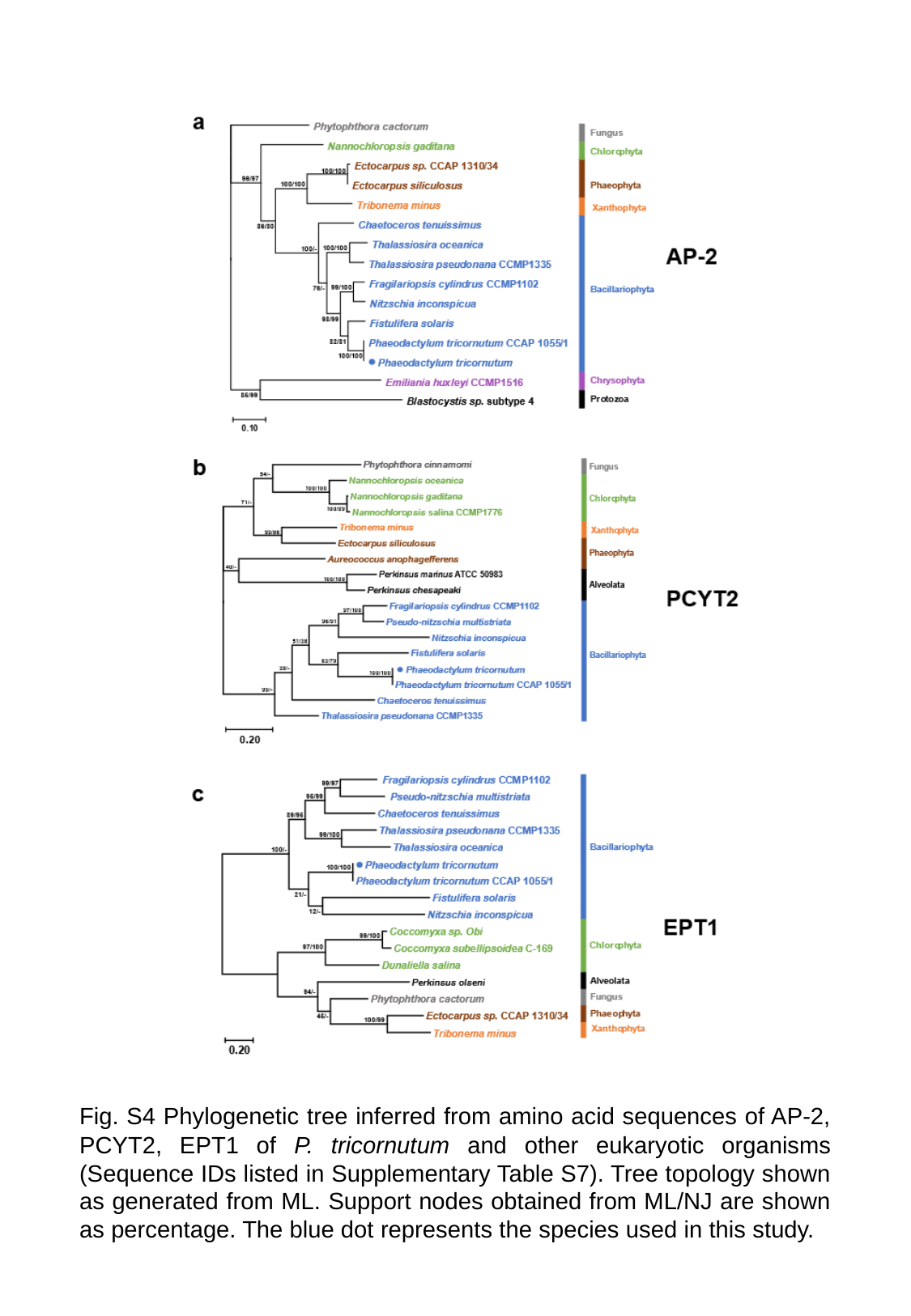

Fig. S4 Phylogenetic tree inferred from amino acid sequences of AP-2, PCYT2, EPT1 of P. tricornutum and other eukaryotic organisms (Sequence IDs listed in Supplementary Table S7). Tree topology shown as generated from ML. Support nodes obtained from ML/NJ are shown as percentage. The blue dot represents the species used in this study.

### Slide 4
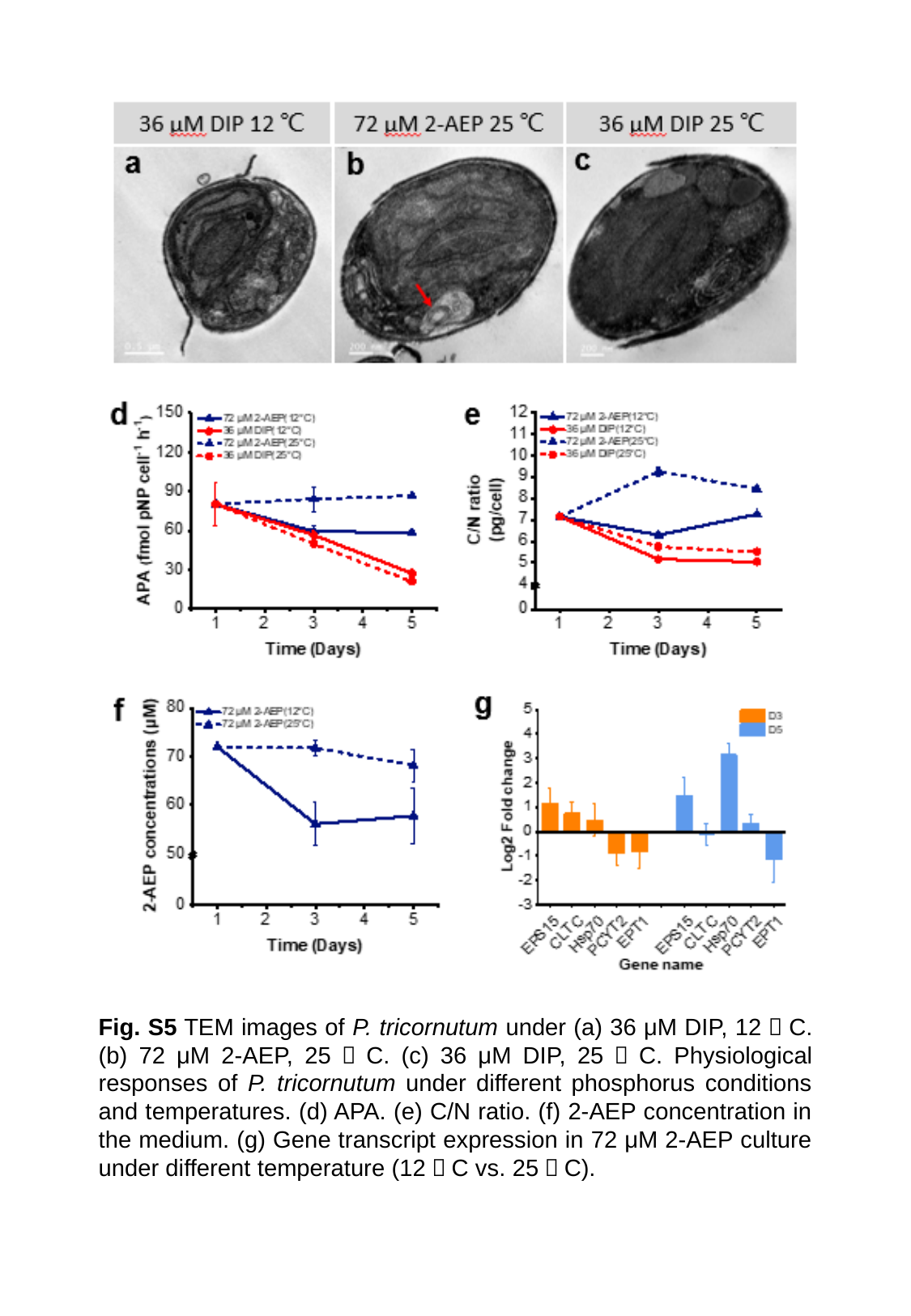

Fig. S5 TEM images of P. tricornutum under (a) 36 μM DIP, 12゜C. (b) 72 μM 2-AEP, 25゜C. (c) 36 μM DIP, 25゜C. Physiological responses of P. tricornutum under different phosphorus conditions and temperatures. (d) APA. (e) C/N ratio. (f) 2-AEP concentration in the medium. (g) Gene transcript expression in 72 μM 2-AEP culture under different temperature (12゜C vs. 25゜C).

### Slide 5
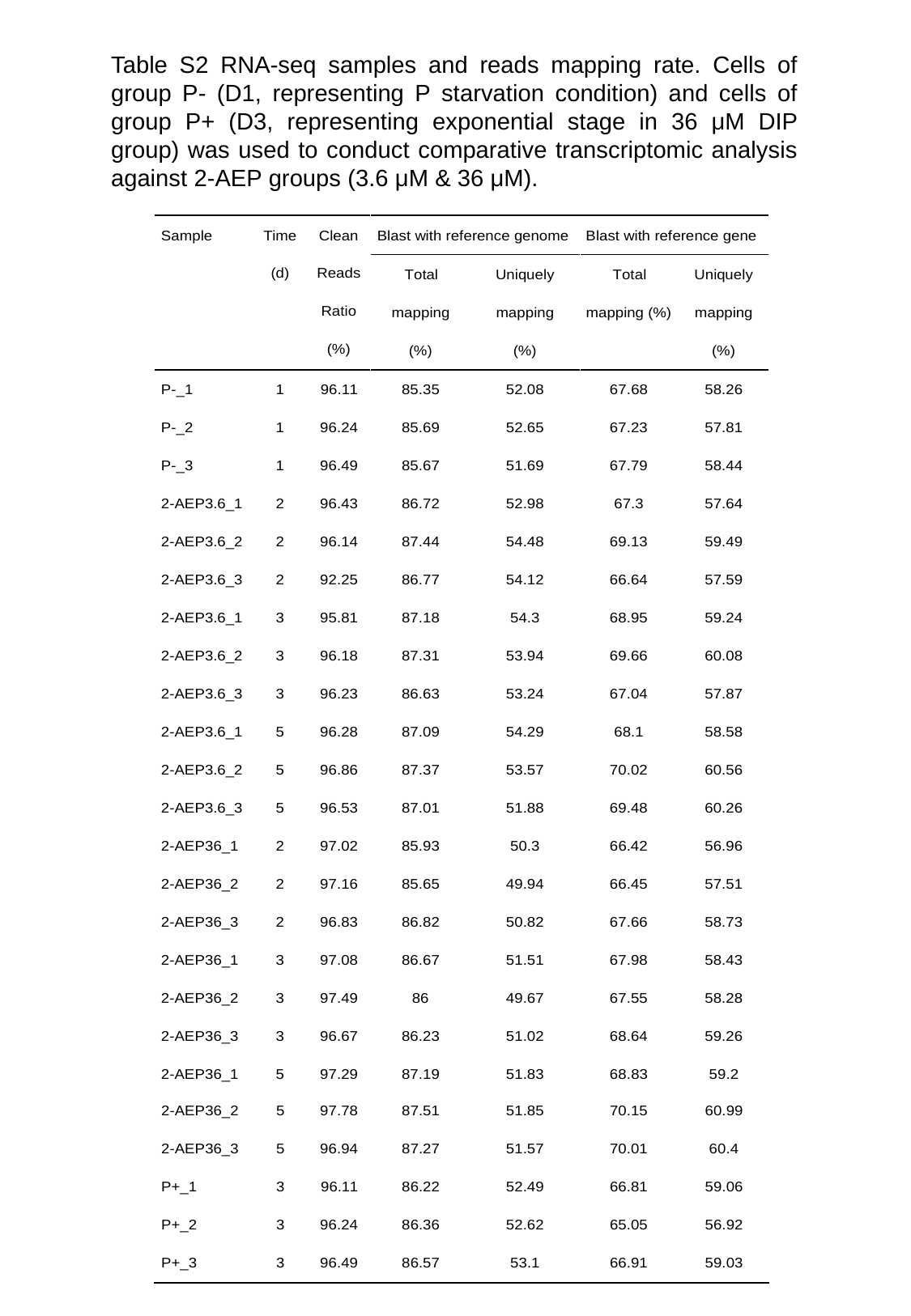

Table S2 RNA-seq samples and reads mapping rate. Cells of group P- (D1, representing P starvation condition) and cells of group P+ (D3, representing exponential stage in 36 μM DIP group) was used to conduct comparative transcriptomic analysis against 2-AEP groups (3.6 μM & 36 μM).

### Slide 6
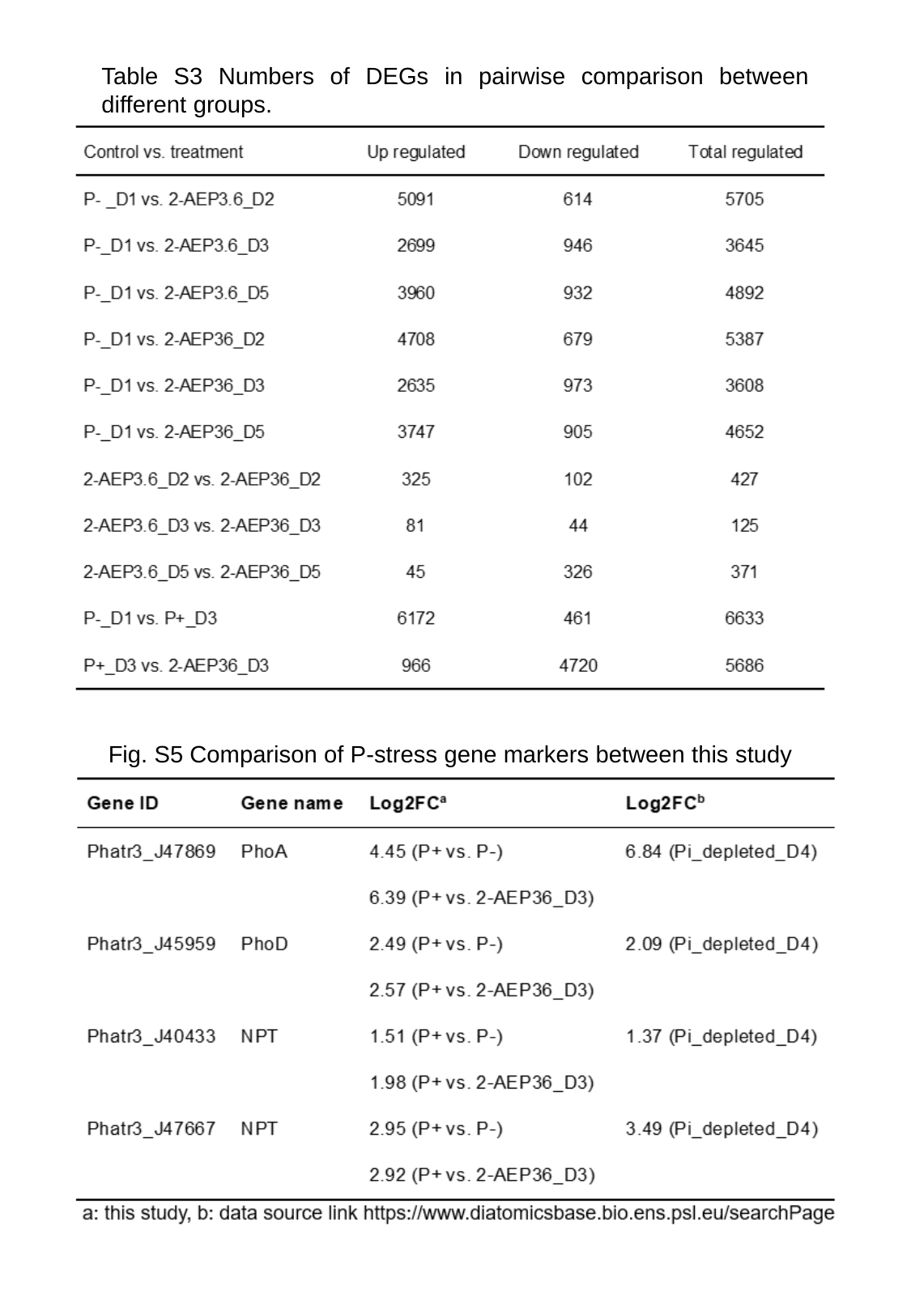

Table S3 Numbers of DEGs in pairwise comparison between different groups.
 Fig. S5 Comparison of P-stress gene markers between this study

### Slide 7
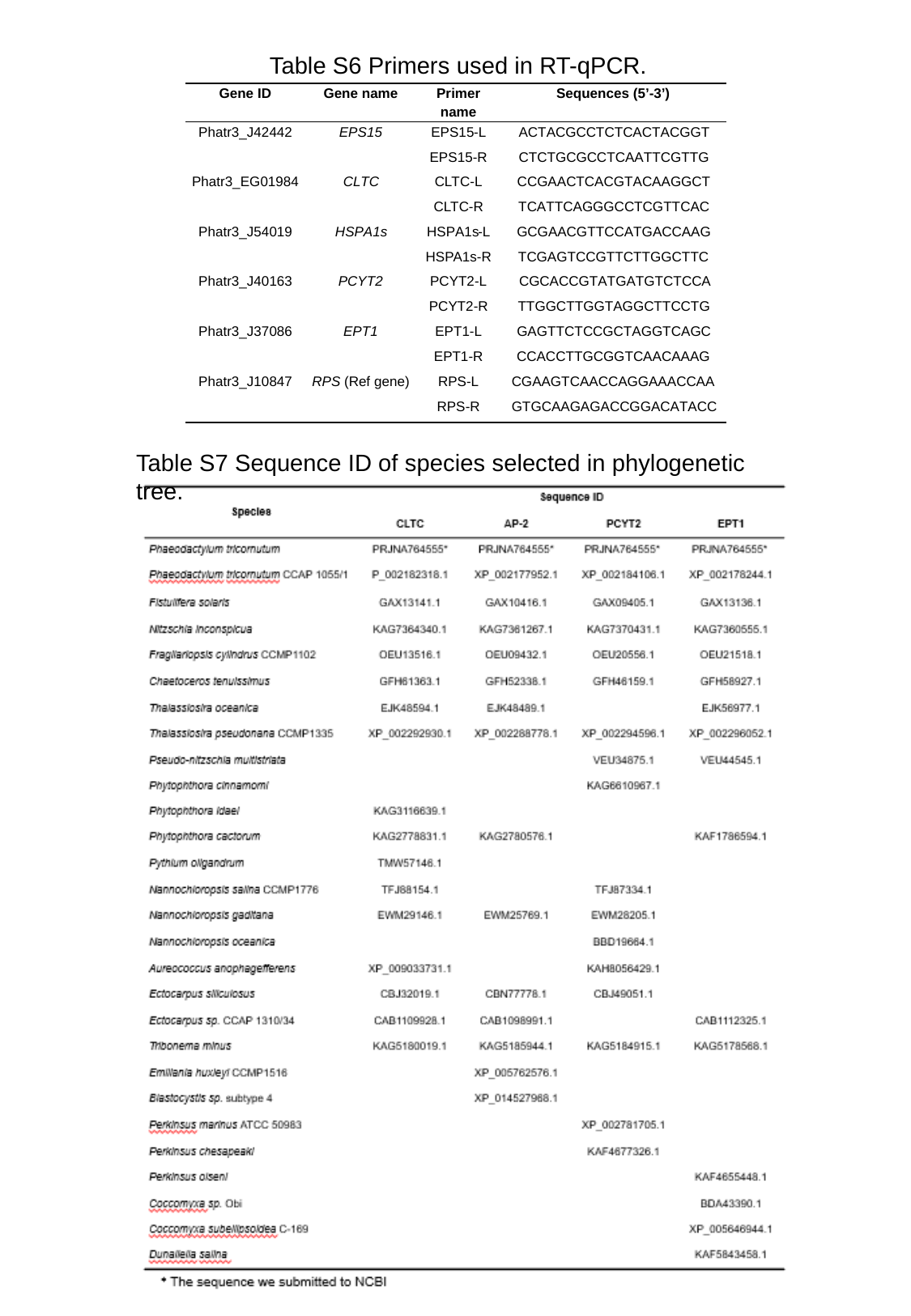

Table S6 Primers used in RT-qPCR.
Table S7 Sequence ID of species selected in phylogenetic tree.
